# MiR-125b differentially impacts mineralization in dexamethasone and calcium-treated human mesenchymal stem cells

**DOI:** 10.1101/2024.08.23.609420

**Authors:** Virginie Joris, Elizabeth R. Balmayor, Martijn van Griensven

## Abstract

Bone metabolism is highly regulated, and microRNAs (miR) can contribute to this process. Among them, miR-125b is well-known to enhance osteoporosis and reduce osteogenic differentiation of human mesenchymal stem cells (hMSCs). In this work we aim to evaluate and understand how miR-125b modulates mineralization of hMSCs in two different *in vitro* models. Cells were cultured in dexamethasone or calcium medium and transfected with miR-125b mimic. Exposure to dexamethasone or calcium medium increased mineralization of hMSCs and was associated with decreased miR-125b expression. Transfection of miR-125b mimic in dexamethasone-treated cells increased mineralization while it decreased it in calcium-treated cells. Levels of osteogenic markers presented the same difference. We identified STAT3, p53 and RUNX2 as direct targets of miR-125b in hMSCs. While these targets remained identical in both treatments, their modulation after transfection was different. We showed that miR-125b mimicking differentially modulated the expression of the miR-199a/214 cluster, probably via STAT3/miR-199a/214, and p53/miR-214 pathways. In conclusion, miR-125b affinity for targets implicated in bone remodeling changed depending on the *in vitro* models used to induce mineralization and led to opposite physiological effect. This works shows the complexity of drugs such as dexamethasone and opens the door for new *in vitro* models of mineralization.

## Introduction

Osteoporosis is a silent systemic skeletal disorder characterized by deregulations of bone homeostasis leading to the deterioration of bone tissue and bone fragility^1^. Recent estimations showed that 33% of women and 20% of men are at risk of developing osteoporosis ^1^, making it a burdening health problem. Disruption of bone homeostasis during osteoporosis is happening through an increase of bone resorption, mediated by osteoclasts and a decrease of bone formation mediated by osteoblasts. While strategies targeting osteoclast activity already showed promising results in the clinic to halt or reverse osteoporosis, stimulating osteoblasts maturation to induce bone formation is a less developed approach.^2^ Moreover, mesenchymal stem cells (MSCs) appear to be an important component to maintain the physiological balance between bone resorption and formation. Indeed, these cells can differentiate into several other cell types as chondrocytes, adipocytes, and osteoblasts.^3^

Regarding the differentiation process of MSCs into osteoblasts, several transcription factors are required, comprising Runt-related transcription factor 2 (RUNX2), Osterix, and β-catenin.^4, 5^ The expression profile of RUNX2 is strictly regulated as it presents a high level in preosteoblasts and then reduces to allow osteoblast maturation and differentiation into osteocytes, showing the dual role of this transcriptional factor in bone remodeling.^5, 6^ Pathways implicated in the differentiation of MSCs into osteoblasts are already well described and comprise important axes such as bone morphogenetic protein (BMP), Wingless/Integrated (WNT), transforming growth factor (TGF)-β, or fibroblast growth factor (FGF) pathways, all leading to higher levels or activity of RUNX2.^4^ Interestingly, in the elderly, more prompt to osteoporosis, the differentiation potential of MSCs into osteoblast decreases in favor of adipocyte differentiation.

The strict and fine-tuned regulation of the pathways implicated in bone remodeling and diseases comprises small biological molecules called microRNAs (miRNAs). MiRNAs are small, highly conserved, non-coding RNAs involved in post-transcriptional regulation through the targeting of the 3’ untranslated region (UTR) of messenger RNAs (mRNAs), and degradation of the targeted mRNA or inhibition of their translation. Over the past years, miRNAs appeared as being able to modulate bone homeostasis.^7^ The conditional skeletal gene–mediated inactivation of *Dicer*, the ribonuclease responsible for the maturation of pre-miRNA into miRNA, highlighted the role of miRNAs in both early and late steps of osteogenesis. Indeed, the ablation of *Dicer* in osteoprogenitor cells leads to bone defects in the embryo, highlighting the importance of miRNAs in bone development.^8, 9^ On the contrary, deleting *Dicer* in mature osteoclasts in adult mice leads to higher bone mass.^10^ This observation suggests the existence of miRNAs that are osteo-enhancers while others are osteo-suppressors.

Previous studies showed that specific miRNAs influence bone mineral density.^8^ Moreover, five circulating miRNAs (miR-21, miR-24, miR-23, miR-100-5p, and miR-125b) were reported as overexpressed in the plasma and at the fracture site of osteoporotic patients compared to healthy.^11,12^ Among these miRNAs, miR-125b is described as able to modulate bone-related pathways.^8, 13, 14^

While its role in bone remodeling is not totally understood, several studies showed miR-125b is able to reduce BMP2 and BMP4-induced differentiation of hMSCs.^15^ Chen and colleagues highlighted an overexpression of miR-125b in osteoporotic bone marrow-derived mesenchymal stem cells (BMSC) compared to healthy patients^16^. They also showed that an overexpression of miR-125b in hBMSCs inhibits their osteogenic differentiation.^16^ Moreover, this miRNA is also able to regulate osteogenesis by targeting cJun, BMP2, or BMPR1b.^14^ Interestingly, miR-125b is also implicated in inflammation through the downregulation of TRAF6 and A20, increasing p65, IL-1β, and IL-8 levels in HT29 cells.^17^ It is therefore not surprising that this miRNA is a focus of interest in bone diseases and remodeling.

*In vitro,* dexamethasone has been used as the gold standard model used to induce differentiation and mineralization of hMSCs^18^. However, the use of this glucocorticoid can be controversial, as, *in vivo*, it induces bone loss^19^. In this work, we aimed to investigate the impact of miR-125b-5p modulation in dexamethasone and calcium-induced mineralization of hMSCs. We observed that miR-125b induced more mineralization in dexamethasone-treated cells while it reduced it in calcium-treated cells. We therefore deepened our investigation to understand the mechanism behind these opposite effects.

## Results

### Mimicking miR-125b-5p induced an opposite effect in dexamethasone and calcium-treated hMSCs

Mineralization of hMSCs was induced using dexamethasone or calcium medium and transfection was performed during the process as described in Fig. 1A and B. After 21 days of culture, hMSCs that were stimulated with dexamethasone medium mineralized while control cells did not (Fig.2A and B). Interestingly, the transfection of miR-125b-5p mimic in dexamethasone-stimulated cells induced a higher mineralization as visualized with Alizarin Red (Fig.2C) and quantified by cetylpyridinium chloride (CPC) method (Fig.2D, p-value<0.001). Expression of miR-125b-5p was assessed by qPCR and revealed a significant decrease of its expression in dexamethasone-treated cells compared to control while after transfection with its mimic, we observed a 11-fold increase of miR-125b-5p expression (Fig.2E, p-value<0.01). In a calcium-saturated medium, after 14 days of culture, hMSCs transfected with a negative sequence (scramble) presented higher mineralization than control (Fig.2F and G). Moreover, in calcium-stimulated cells, the transfection of mimic 125b-5p led to a decrease in mineralization (Fig.2H) as also quantified by CPC method (Fig.2I, p-value<0.01). Similar to what was observed for dexamethasone-containing medium, miR-125b-5p expression was significantly downregulated in calcium-saturated medium and the mimic transfection induced a 7.1-fold upregulation of its expression (Fig.2J, p-value<0.05).

**Figure 1:**
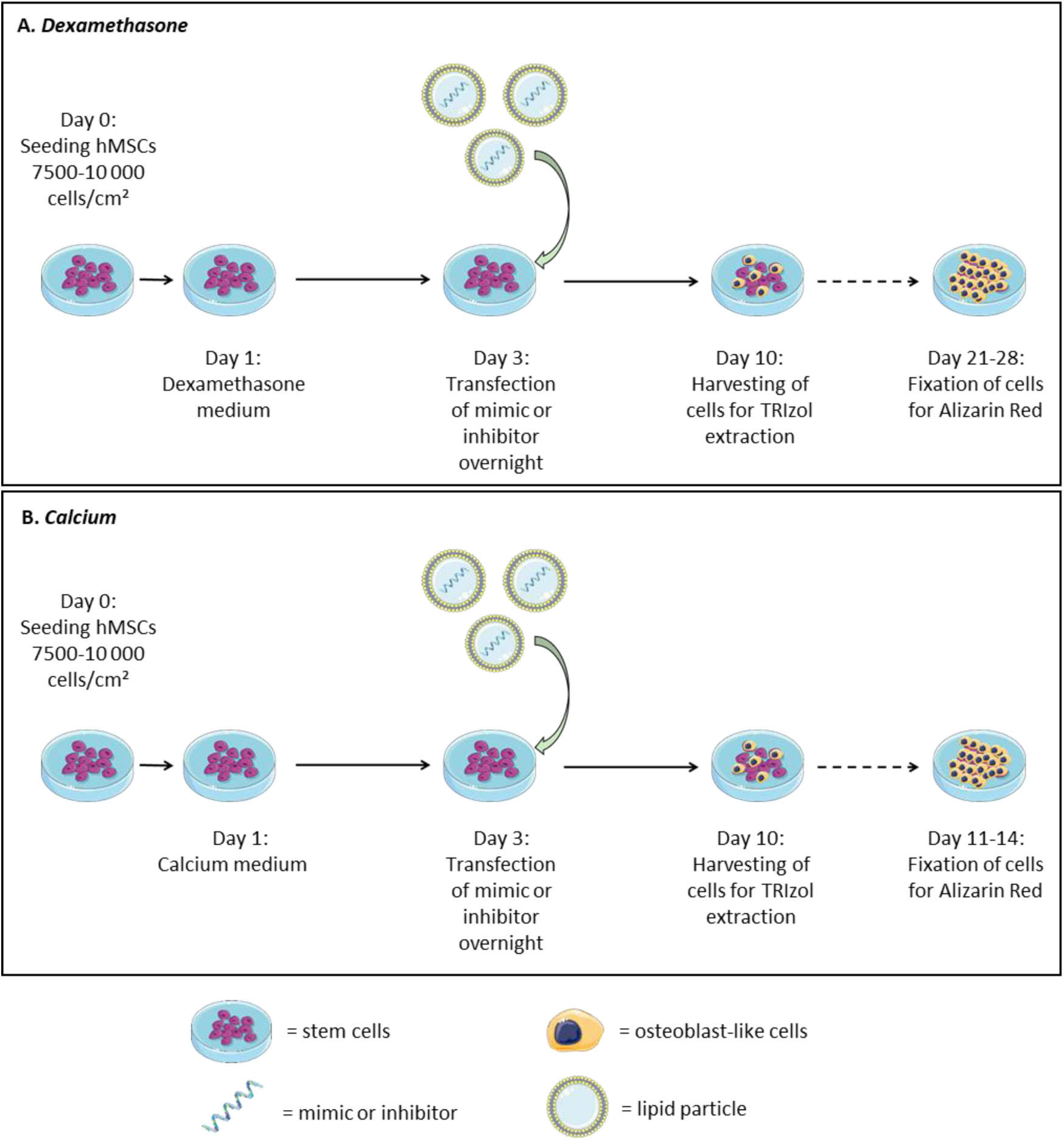
Schematic representation of dexamethasone and calcium-induced mineralization. Dexamethasone (A) or calcium chloride (B) was added to the cells one day after seeding. Transfection with mimics or negative control was performed at day 3 and cells were harvested 10 days after the beginning of dexamethasone or calcium treatment. Mineralization was assessed via Alizarin Red after 21-28 days for dexamethasone and 11-14 days for calcium protocol. Dexamethasone medium = dexamethasone 100 nM, beta-glycerol phosphate and ascorbic acid. Calcium medium = Calcium chloride 8 mM and ascorbic acid.

**Figure 2:**
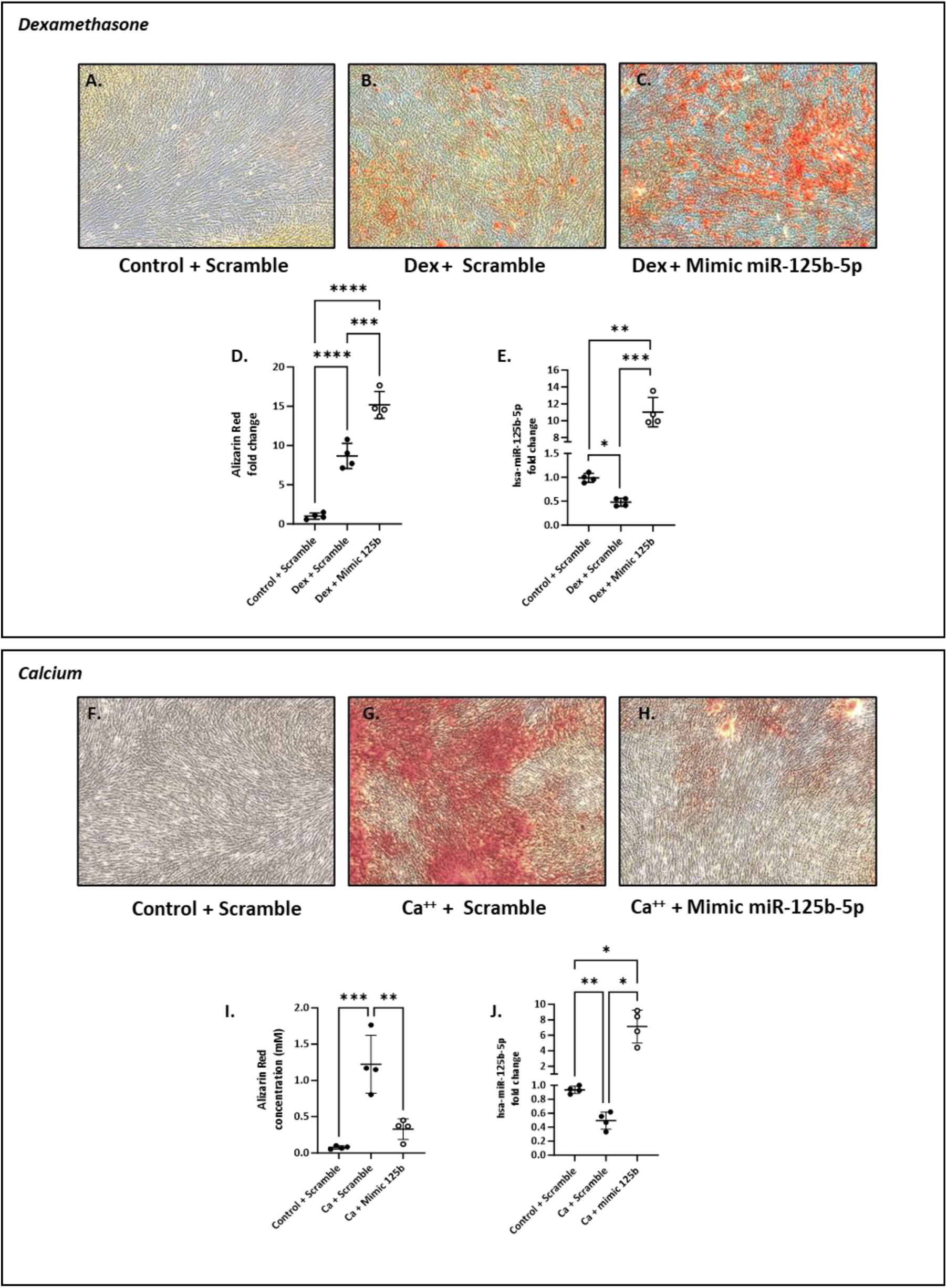
Mimicking of miR-125b induced opposite effect in dexamethasone and calcium-treated hMSCs. hMSCs were transfected with negative control or mimic and cultured in control or dexamethasone medium (upper panel) and in control or calcium medium (lower panel). Mineralization was assessed with alizarin red ( A-C and F-H) and quantified with CPC (D and I) using a spectrophotometer, \*\**p* < 0,01; \*\*\**p* < 0.001; \*\*\*\**p* < 0.0001. miRNA-125b expression was assessed using qPCR (E and J) \**p* < 0.05; \*\**p* < 0.01; \*\*\**p* < 0.001 based on fold change relative to control + scramble.

### BMPR2 and RUNX2 proteins presented a pattern corroborating the mineralization state

After 7 days, BMPR2 and RUNX2 levels were significantly increased in scrambled-transfected hMSCs after dexamethasone stimulation compared to control cells (Fig3.A, B and C). Similarly to what we observed with Alizarin red staining, their level was even higher in the dexamethasone-treated group transfected with the miR-125b-5p mimic. Moreover, *USP7* expression, a stabilizer of RUNX2, was increased with dexamethasone and remained upregulated when the miR-125b mimic was transfected (Fig.3D, p-value<0.001). Interestingly, we showed that RUNX2 is a direct target of miR-125b in dexamethasone-treated cells (Fig.3E). HMSCs transfected with scrambled mimic and cultured in the presence of calcium also presented increased levels of BMPR2 and RUNX2 compared to control cells.

**Figure 3:**
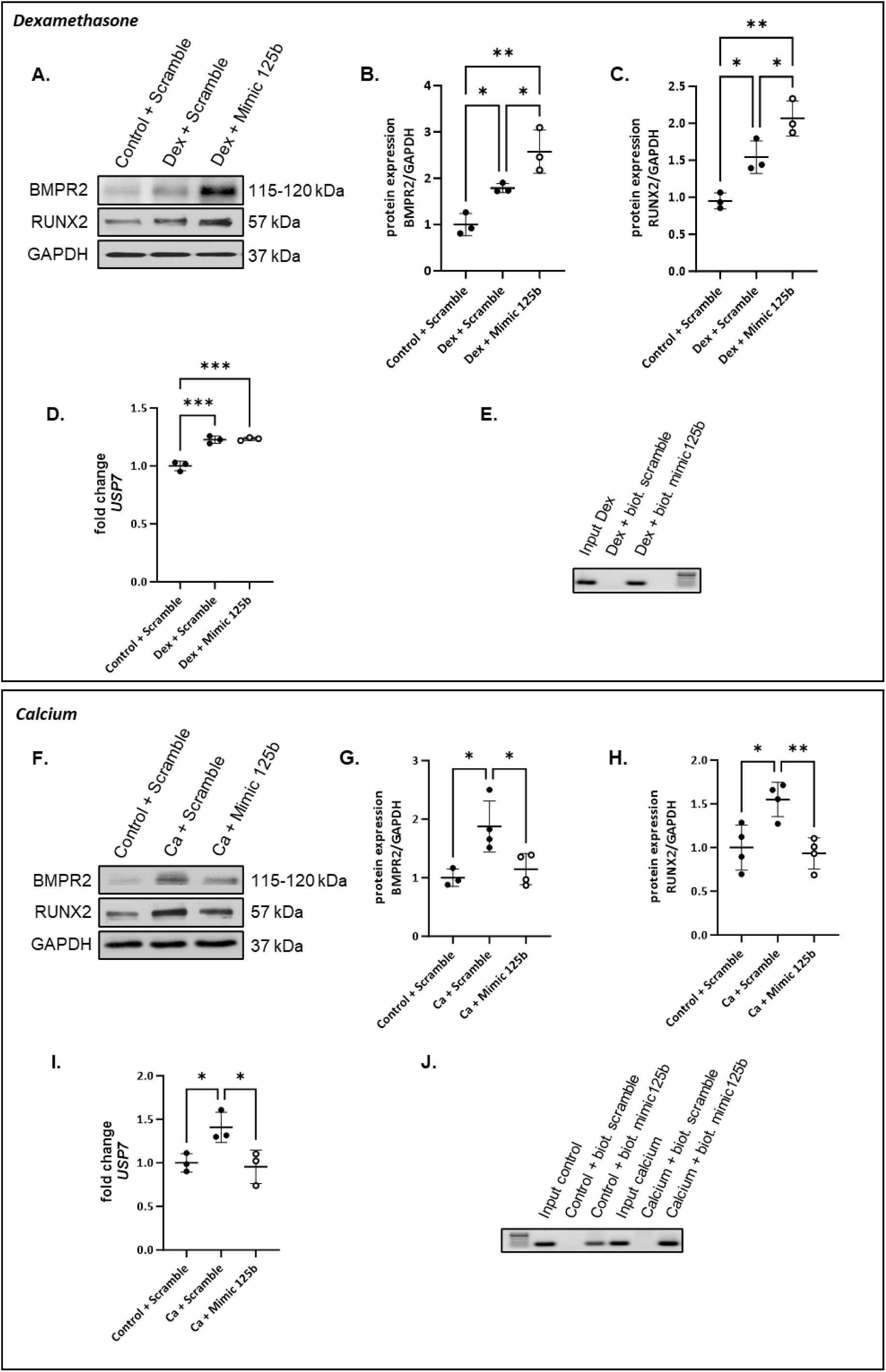
Mimicking of miR-125b induced the opposite profile of osteogenic proteins level. hMSCs were transfected with negative control or mimic and cultured in control or dexamethasone medium (upper panel) and in control or calcium medium (lower panel). BMPR2 and RUNX2 levels were measured via western blotting (A and F). Quantification of protein levels was performed using ImageJ normalized to GAPDH (B-C and G-H) \**p* < 0.05; \*\**p* < 0.01. *USP7* expression was assessed using qPCR (D and I) \**p* < 0.05; \*\*\**p* < 0.001 based on fold change relative to control + scramble. The verification of RUNX2 as a direct target of miR-125b was performed using a pull-down assay with biotinylated mimic-125b and negative control combined with PCR and agarose gel (E and J).

However, when transfected in calcium-treated cells, the miR-125b-5p mimic induced a significant decrease of these two proteins (Fig.3F, G and H). While *USP7* expression is increased in calcium-treated cells, it returns to control levels in cells transfected with the mimic (Fig.3I, p-value<0.05). As observed for dexamethasone-treated cells, RUNX2 was observed to be a direct target of miR-125b in calcium-treated cells (Fig.3J).

### Mimicking miR-125b-5p differentially regulated miR-199a/214 cluster in dexamethasone and calcium-treated hMSCs

The expression of miR-199a-5p and miR-214-3p was measured in hMSCs stimulated with dexamethasone or calcium for 7 days and transfected with scramble or miR-125b-5p mimic. We observed that both dexamethasone (Fig.4A and B) and calcium (Fig.4C and D) treatments induced a significant downregulation of miR-199a-5p and miR-214-3p in hMSCs. However, transfection of miR-125b-5p mimic in dexamethasone-stimulated cells emphasized this downregulation (Fig.4A and B, p-value<0.05 and 0,01) while it counteracted it in calcium-stimulated cells. Indeed, after transfection with miR-125b-5p mimic, calcium-treated cells presented a higher expression of the miR-199a/214 cluster compared to calcium-treated cells transfected with the scrambled-miR (Fig.4C and D, p-value<0.01). Interestingly, in dexamethasone-treated cells, the downregulation of miR-199a/214 cluster is associated with a significant increase of caveolin-1, a well-known target of miR-199a-5p (Fig.S1).

**Figure 4:**
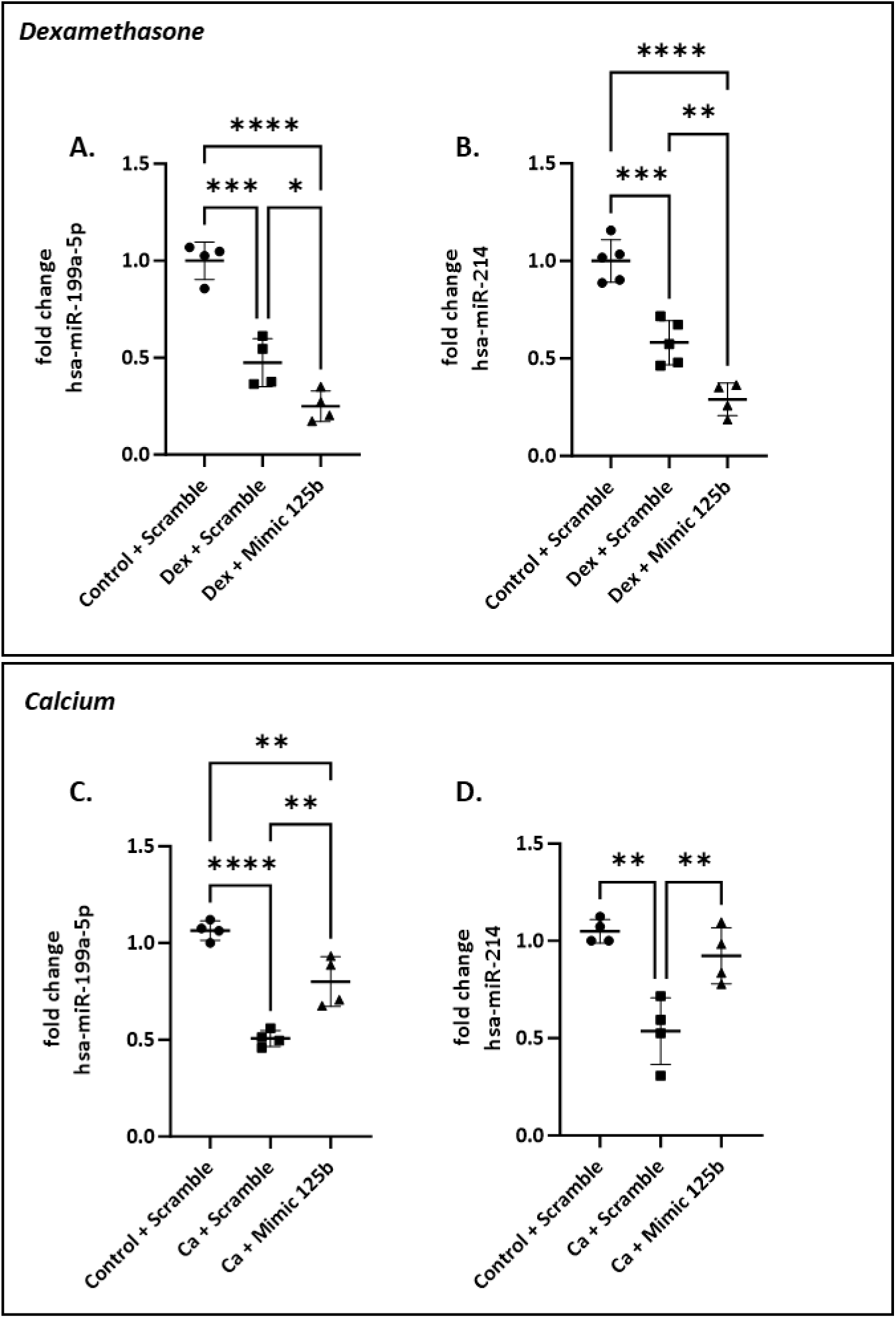
Mimicking miR-125b-5p differentially regulated miR-199a/214 cluster in dexamethasone and calcium-treated hMSCs. hMSCs were transfected with negative control or mimic and cultured in control or dexamethasone medium (upper panel) and in control or calcium medium (lower panel). MiR-199a-5p and miR-214-3p expression was measured using qPCR. \**p* < 0.05; \*\**p* < 0.01; \*\*\**p* < 0.001; \*\*\*\**p* < 0.0001 based on fold change relative to control + scramble.

### Signal transducer and activator of transcription 3 (STAT3) and Tumor protein p53 were direct targets of miR-125b in hMSCs but behaved differently in dexamethasone and calcium-treated cells transfected with mimic

After performing a pull-down assay using biotinylated miR-125b mimic or negative control, we observed that *STAT3* and *p53* were direct targets of miR-125b (Fig. 5A). However, their expression did not vary the same in transfected dexamethasone or calcium-stimulated cells. Indeed, while transfection of miR-125b in dexamethasone-treated cells did not induce any modification of *STAT3* expression (Fig. 5B), *p53* expression decreased compared to other conditions (Fig. 5C, p-value<0.001). At the opposite, calcium treatment induced a significant overexpression of *STAT3* and the concomitant transfection of miR-125b counteracted it (Fig. 5D, p-value<0.01) while *p53* expression was not modified (Fig. 5E)

**Figure 5:**
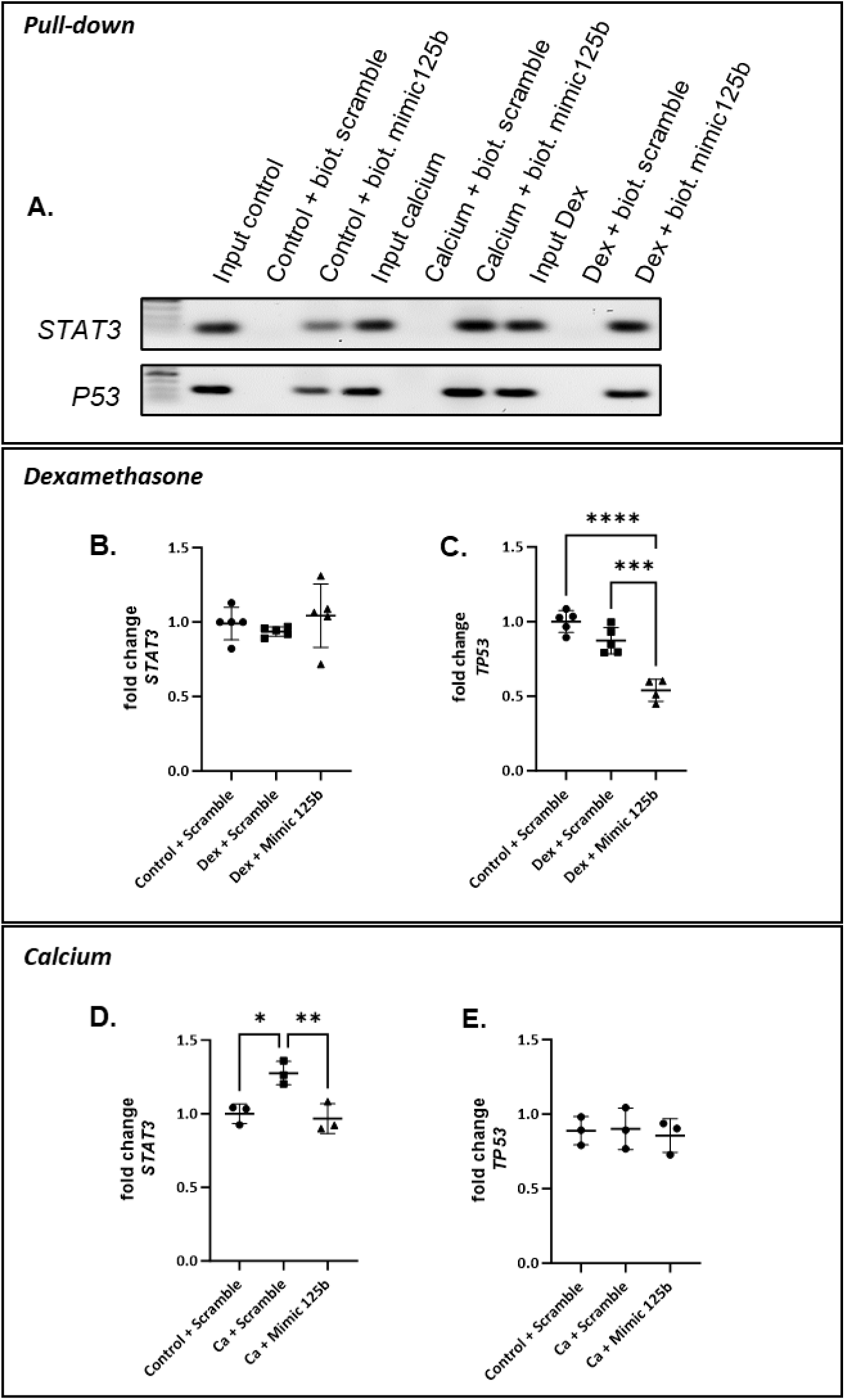
The direct targets STAT3 and p53, reacted differently to miR-125b in dexamethasone and calcium medium. The verification of STAT3 and TP53 as a direct target of miR-125b was performed using a pull-down assay with biotinylated mimic-125b and negative control combined with PCR and agarose gel (A). hMSCs were transfected with negative control or mimic and cultured in control or dexamethasone medium (middle panel) and in control or calcium medium (lower panel). STAT3 (B and D) and p53 (C and E) expression was measured using qPCR. \**p* < 0.05; \*\**p* < 0.01; \*\*\**p* < 0.001; \*\*\*\**p* < 0.0001 based on fold change relative to control + scramble.

### Rescuing miR-214 expression in dexamethasone-treated hMSCs transfected with miR-125b mimic abrogated mineralization

To understand if miR-214 could be the culprit of the effect observed in dexamethasone-treated hMSCs, we co-transfected miR-125b and miR-214 mimics. We observed that, as previously reported, dexamethasone induced mineralization of hMSCs compared to control medium (Fig 6. A and B). However, while the transfection with miR-125b mimic enhanced the mineralization induced by dexamethasone (Fig. 6C), the cells transfected with miR-125b and miR-214 mimics did not show mineralization *per se* (Fig. 6D). Measurement of miR-125b and 214 expressions confirmed the efficiency of the transfection (Fig. 6 E and F).

**Figure 6:**
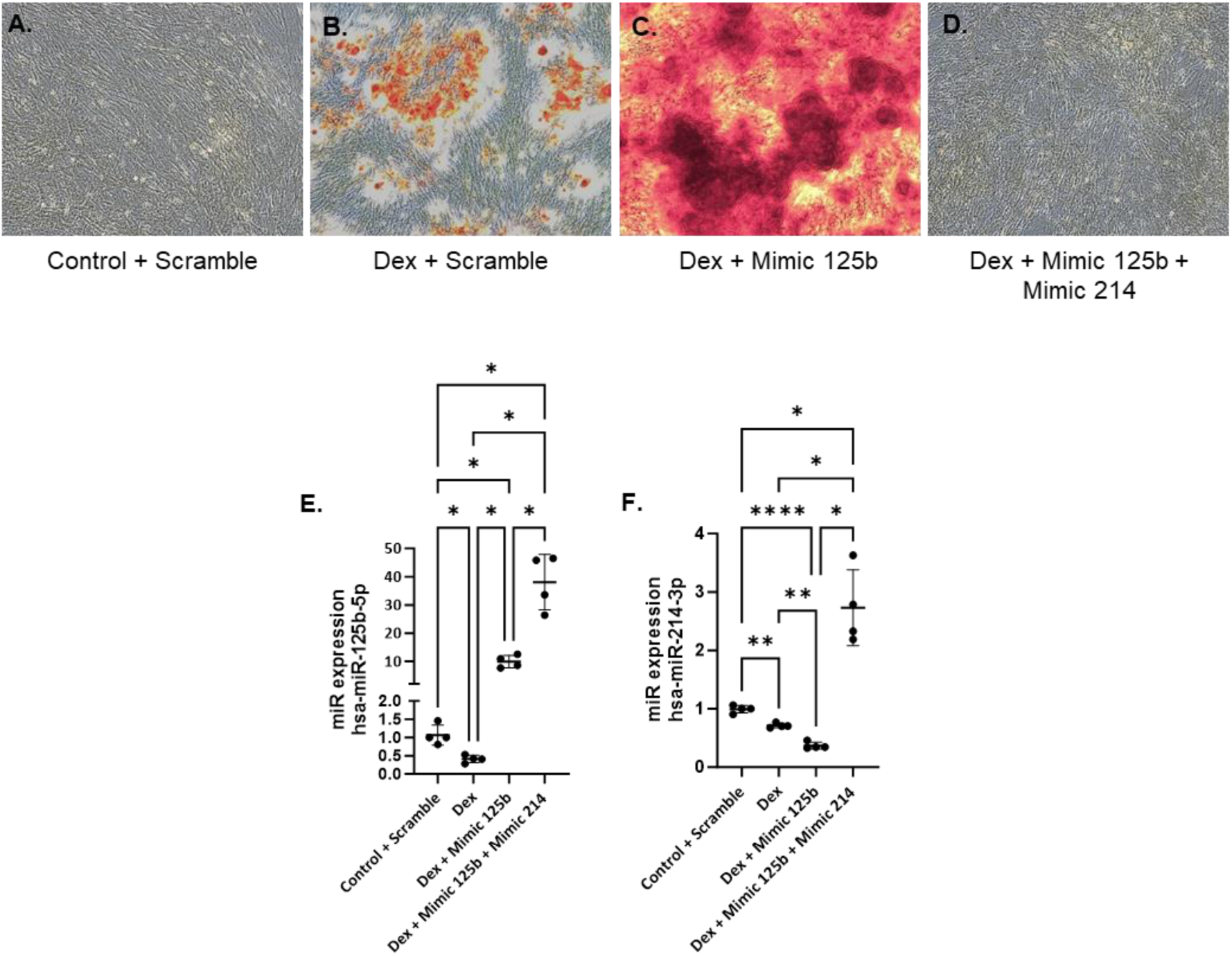
Rescuing miR-214 in dexamethasone-treated hMSCs transfected with miR-125b abrogated mineralization. hMSCs were transfected with negative control or mimic and cultured in control or dexamethasone medium. Mineralization was assessed with alizarin red (A-D). miRNA-125b and miR-214-3p expression was assessed using qPCR (E and F). \**p* < 0.05; \*\**p* < 0.01; \*\*\*\**p* < 0.0001 based on fold change relative to control + scramble.

## Discussion

Musculoskeletal disorders are considered a burdening health problem with osteoporotic fractures as a major contributor. Osteoporosis is a silent disease until a fracture occurs and, for now, the available treatments are causing several adverse effects. Osteoporosis is characterized by an imbalance between bone formation and resorption, leading to a more fragile bone.^1^ The mechanisms behind bone remodeling are partially known and require several pathways as BMP, FGF, TGF and Wnt.^4^ However, fine-tuned regulation of these pathways is not yet well studied and understood.

In this work, we tried to elucidate the role of miR-125b in bone metabolism. Indeed, miR-125b is a member of a five miRNAs signature found in the blood and fracture site of osteoporotic patients.^11,12^ Here, we showed that the modulation of miR-125b in hMSCs induced different effects on mineralization depending on the *in vitro* model used. We observed an increase of mineralization in dexamethasone-treated hMSCs with miR-125b. However, we were surprised to find that the concomitant mimicking of miR-125b in those cells induced a higher mineralization than the dexamethasone alone. In fact, several studies reported miR-125b to be able to reduce osteogenic differentiation of MSCs. Mizuno *et al.* showed that overexpression of miR-125b induces the inhibition of proliferation of ST2 cells, followed by the suppression of BMP4-induced osteogenesis.^20^ Huang *et al.* reported that miR-125b expression is decreased during osteogenic differentiation of C3H10T1/2 cells, and its overexpression reduces mineralization through downregulation of Cbfβ gene.^15^ In bone marrow-derived MSCs, the overexpression of miR-125b leads to the suppression of osteogenic differentiation, probably by targeting osterix.^16^ Wang *et al*. showed that, in hMSCs, miR-125b can reduce osteogenic differentiation via a downregulation of BMPR1b.^14^ *In vivo*, the role of miR-125b in bone is more shaded. While some studies show that miR-125b overexpression blocks osteoclastogenesis and promotes higher bone mass^21^, other works design miR-125b as a culprit of osteoporotic phenotype.^13^

To further investigate the results obtained, we wanted to reproduce it in another osteogenic *in vitro* model consisting of calcium-saturated medium using calcium chloride. The use of extracellular calcium ions to induce mineralization of MSCs is already well developed using both calcium-coated surface or calcium ion-saturated medium.^22–24^ In this model, we observed that culturing the cells in calcium medium increased the mineralization of hMSCs, as observed for dexamethasone. However, in this condition, the mimicking of miR-125b drastically reduced this effect. Interestingly, the difference observed in mineralization between dexamethasone and calcium-treated cells was associated with an opposite pattern of osteogenic proteins such as BMPR2 and RUNX2. Indeed, while both treatments induced an increase of the two proteins, miR-125b mimic emphasized the upregulation in dexamethasone-treated cells and counteracted it in calcium-treated cells. While BMPR2 is a well-known target of miR-125b in musculoskeletal disorders ^13, 25^, RUNX2 is always described as an indirect target.^26^ In this work, we showed, for the first time to our knowledge, that RUNX2 is a direct target of miR-125b in hMSCs. Moreover, we also observed a differential expression of USP7, known to deubiquitinate RUNX2 and avoid its degradation by the proteasome. Kim *et al.* showed that knockdown of USP7 in MSCs reduce RUNX2 and osteopontin levels while another study highlighted the importance of deubiquitination of RUNX2 by USP7 for bone formation.^27, 28^ These results indicate that, in our model, miR-125b might be able to impact RUNX2 by regulating both its expression and stabilization.

To better understand the reason for the opposite effect obtained in dexamethasone and calcium-treated hMSCs transfected with miR-125b mimic, we performed *in silico* studies. We highlighted two potential targets: STAT3 and p53. In their study about osteosarcoma, Liu *et al*. showed that miR-125b can directly target STAT3 thereby reducing proliferation and migration of the cancerous cells.^29^ Moreover, miR-125b is also described as a negative regulator of p53 in neuroblastoma cells where it reduces their apoptosis.^30^ More recently, a study showed that increased levels of miR-125b in the human breast cancer cell line MCF7 induced a decrease of p53, and this was associated with DNA damage.^31^ In the hMSCs used in this study, we confirmed, via pull-down assay, that STAT3 and p53 are targets of miR-125b in both dexamethasone and calcium-treated cells. However, their expression showed a different pattern in both treatments. Interestingly, calcium is known to induce STAT3 expression, while dexamethasone alone is not.^32, 33^ This corroborates what we observed in hMSCs. Opposite to our observations, dexamethasone was previously reported to increase p53 in MC3T3-E1 osteoblastic cells.^34^ However, this has been mostly investigated at the protein level and not RNA level, as investigated in the present study.

The identification of STAT3 and p53 as direct targets of miR-125b led us to a specific cluster of miRNAs, miR-199a/214 cluster. Indeed, STAT3 is reported as a modulator of miR-199a and miR-214 expression in cardiac and thyroid tissues.^35, 36^ P53 is also associated with miR-199a and miR-214 expression during renal fibrosis and it has been identified as a transcription factor of these two miRNAs.^37, 38^ With this in mind, we measured the expression of miR-199a-5p and miR-214 and showed that it was also differentially regulated in dexamethasone and calcium-treated cells. Both treatments induced a downregulation of the two miRNAs but mimicking miR-125b in dexamethasone reduced even more their expression. Conversely, overexpression of miR-125b in calcium-treated cells restored the level of miR-199a-5p and miR-214 in hMSCs. Moreover, the rescue of miR-214 in dexamethasone-treated cells transfected with miR-125b mimic drastically reduced the mineralization of the cells. MiR-214 is reported as a biomarker of osteoporosis and it is known that upon production by osteoclasts, this miRNA can inhibit osteoblast differentiation in ovariectomized mice.^39^ Also, Yuan *et al.* showed that shear stress due to physical exercise downregulates the expression of miR-214 in tibial bone and increased expression of osteogenic markers.^40^ While the effect of miR-199a-5p on bone formation is not clear, its downregulation in dexamethasone treated cells was associated with an increase of caveolin1 expression, one of its targets.^41^ Interestingly, BMPR2 is localized in caveolae and the downregulation of caveolin1 leads to reduced localization of BMPR2 at the membrane.^42^ Therefore, the upregulation of caveolin1 observed in dexamethasone-treated cells could contribute to the increased mineralization by increasing the amount of BMPR2 at the basal membrane.

Dexamethasone is often used to induce the differentiation of stem cells into osteogenic lineages.^43^ However, its effect on osteogenic differentiation is controversial. Prolonged use of dexamethasone *in vivo* induces osteoporosis and increases the risk of fracture.^19^ Moreover, Li *et al.* showed that BMSCs isolated from dexamethasone-treated mice differentiate into adipocytes instead of osteoblasts.^44^ Already twenty years ago, a study showed that dexamethasone can inhibit the mineralization of osteoblasts in a dose and time-dependent manner.^45^ Nowadays the controversy around dexamethasone is not yet elucidated. Recently, dexamethasone was shown as able to induce osteogenic differentiation of MSCs through downregulation of SOX9 and not through upregulation of RUNX2 as usually thought.^46^ This was also correlated with an upregulation of PPARγ, inducing the appearance of adipocyte-like cells and showing that dexamethasone is probably not the best *in vitro* model to induce osteogenesis.^46^ Our present work aligns with this last statement. Results obtained when transfecting dexamethasone-treated cells with miR-125b mimic align with what is observed in the literature. Furthermore, our obtained results support the statement that miRNA expression profile is dependent on the specific in vitro mineralization model used.

In conclusion, in this work, we showed that dexamethasone and calcium *in vitro* mineralization models react differently to overexpression of miR-125b. We showed that while the targets of miR-125b are the same in the two treatments, their regulation by the miR is different and impact the downstream cascade including miR-199a/214 cluster. To our knowledge, this is the first time that the overexpression of miR-125b is reported as inducing a modulation of other miRNAs. Furthermore, the treatment-dependent difference reported in this work emphasizes the importance of *in vivo* or clinical study design. Indeed, treatments such as dexamethasone are useful to treat certain pathologies but remain highly stressful and, as observed in this study, may deeply impact the cellular response, even at the epigenetic level. To better understand the mechanisms behind the differential response observed in the two treatments, a more detailed study must be performed focusing on miR-199a and 214. This work also draw attention to miR-125b, 119a-5p and 214 as new potential therapeutic targets for bone diseases.

## Materials and Methods

### Culture of human mesenchymal stem cells (hMSC)

Primary hMSCs were isolated from bone marrow provided by the department of trauma surgery from the MUMC+ university hospital from at least 5 different donors (mean age, 2 males, 2 females, 1 unknown). Informed consent was obtained from each donor, as well as approval from the ethical committee (ethical number METC 15-4-274), following the Helsinki guidelines. Patients presenting autoimmune diseases were excluded from the study. Cells were maintained in culture in alpha-minimum essential medium (α-MEM; ThermoFisher Scientific, Waltham, MA, USA) supplemented with 10% fetal bovine serum (FBS; ThermoFisher Scientific, Waltham, MA, USA). The medium was refreshed every two days and cells were passaged using 0.05% trypsin-EDTA (ThermoFisher Scientific, Waltham, MA, USA) when confluency reached 80-90%. For experiments, cells were not used after reaching passage 6. Previous to any experiment, the multipotency of each donor was validated by inducing their differentiation into adipocytes, chondrocytes, or osteoblasts.

### Dexamethasone and Calcium stimulation

HMSCs were seeded at a density of 7500 cells/cm^2^ in a 6-well plate and maintained in α-MEM with 10% FBS. After attachment, cells were cultured in dexamethasone medium consisting of α-MEM with 10% FBS, 10 mM β-glycerophosphate (BGP; Sigma, Saint-Louis, MO, USA), 0.02 mM ascorbic acid phosphate (ASAP), and 100 nM dexamethasone or calcium-saturated medium consisting of α-MEM containing 10% FBS, 8 mM of calcium chloride and 0.02 mM of ASAP (Fig.1). Control cells were maintained in α-MEM supplemented with 10% FBS. After 24h, the cells were transfected overnight (ON) as described below. Subsequently, the cell culture medium was replaced by a differentiation medium. Media changes were performed every two days. For immunoblotting and qPCRs, cells were harvested 7 days after transfection. The alizarin red staining to assess the mineralization of the cells was performed 14 days and 21 days after the induction of calcium or dexamethasone stimulation, respectively.

### Transfection of hMSCs

Transfection of hMSCs was performed 24 h after the beginning of differentiation in both dexamethasone– and calcium-treated cells. Cells were transfected ON with 25 nM of scrambled miR (negative control, Qiagen, Hilden, Germany) or 25 nM of miR-125b-5p mimic (Qiagen, Hilden, Germany) or a combination of 25 nM of miR-125b-5p mimic and 10 nM of miR-214 mimic (Qiagen, Hilden, Germany) using lipofectamine 3000 (Invitrogen, Waltham, MA, USA) in Opti-MEM and diluted in α-MEM with 10% FBS according to the manufacturer’s protocol. Medium was refreshed with basal, osteogenic or mineralization medium and renewed every two days till the end of the culture.

### Extraction of mRNAs and miRNAs

mRNAs and miRNAs were extracted using TRIzol (Invitrogen, Waltham, MA, USA) as advised by the manufacturer’s protocol. Briefly, cells were harvested in TRIzol and chloroform (200 µl/ml TRIzol; Sigma, Saint-Louis, MO, USA) was added. Samples were mixed and incubated at room temperature (RT) for 15 min. After centrifugation for 20 min at 4°C, the aqueous phase was transferred in new tubes and isopropanol (500 µl/ml TRIzol; Fisher Scientific, Hampton, USA) was added, incubated for 15 min at RT, and centrifuged for 20 min at 4°C. Isopropanol was then removed and the pellet was washed twice with a cold 70% ethanol solution. RNA concentration was measured using a Biodrop device (Fisher Scientific, Hampton, NH, USA).

### miRNA reverse transcription and quantitative PCR (qPCR)

Following the manufacturer’s instructions, the first-strand cDNA synthesis was performed using miRCURY LNA RT Kit (Qiagen, Hilden, Germany). Briefly, 40 ng of RNA was loaded in the reaction and a mix containing RT buffer, water, and RT enzyme was added to it to a final volume of 10 µl. Samples were incubated at 42°C for 60 min followed by an inactivation reaction of 5 min at 95°C. Samples were then diluted and 3 µl were loaded in duplicate in a 96-well qPCR plate. The miRCURY SYBR Green Master Mix (Qiagen, Hilden, Germany) along with PCR primers for hsa-miR-125b-5p, hsa-miR-100-5p, hsa-miR-199a-5p, hsa-miR-214-3p or hsa-miR-21-3p were added and PCR run was performed in a CFX96 machine (Bio-Rad, Hercules, CA, USA) following the manufacturer’s protocol. The samples underwent an initial heat activation at 95°C for 2 min followed by a two-step cycling consisting of a first step at 95°C for 10 s and a second at 56°C for 60 s. The two-step cycling was repeated 40 times before melting curve analysis was performed.

### mRNA reverse transcription and qPCR

The first-strand cDNA synthesis was performed using the iScript kit (Bio-Rad, Hercules, CA, USA). 500 ng of RNA were used in the reverse transcription reaction and a mix containing RNAse-free water (Qiagen, Hilden, Germany), RT enzyme, and RT buffer was added to a final volume of 20 µl. Samples were incubated at 25°C for 5 min for a priming step followed by an incubation at 46°C for 20 min and an inactivation step at 95°C for 1 min. cDNA was diluted at a concentration of 5 ng/µl and 2 µl were loaded in duplicate in a 96-well qPCR plate with the reaction mix comprising 5 µl of Sybr Green master mix (Bio-Rad, Hercules, CA, USA), 2 µl of water and 1 µl of primer mix containing forward and reverse primers for *STAT3* and *P53*. The real-time PCR CFX96 was programmed to perform 40 cycles of 10 s of denaturation at 95°C, and 60 s of combined annealing and extension at 60°C. Genes of interest were normalized to the housekeeper gene *GAPDH*.

### Western Blot

Proteins were extracted using TRIzol following the manufacturer’s protocol and quantified using Pierce bicinchoninic acid assay (ThermoFisher Scientific, Waltham, MA, USA). Western blot was performed on an 8%-10% acrylamide gel and 15 µg of proteins were loaded per well. Electrophoresis was performed for 15 min at 90 V in a running buffer consisting of 25 mM Tris, 192 mM glycine, 0.1% SDS, pH 8.3 following dilution to 1x with water and then increased to 120 V for 60 min. After separation, proteins were transferred onto a nitrocellulose membrane with a constant amperage of 350 mA for 90 min in a transfer buffer (50 mM Tris-base, 40 mM glycine, 1.5 mM SDS). Membranes were blocked in TBS-5% bovine serum albumin (BSA)-0.1% Tween-20 for 60 min at RT. Primary antibodies used for immuno-detection were diluted in TBS-5% BSA-0.1% Tween and incubated ON at 4°C. Primary antibodies were: anti-SMAD4 (1/1000; Cell Signaling, Danvers, MA, USA), anti-RUNX2 (1/1000; ThermoFisher Scientific, Waltham, MA, USA), anti-BMPR2 (1/1000, Novus biologicals, Centennial, CO, USA), and anti-GAPDH (1/10000; Cell Signaling, Danvers, MA, USA). Membranes were rinsed with TBS-0.1% Tween-20 and incubated with horse radish peroxidase–conjugated secondary antibodies diluted in TBS-5% BSA-0.1% Tween-20 for 60 min at RT. Development was performed using Bio-Rad Clarity substrate ((Bio-Rad, Hercules, CA, USA) and CL-Xposure film (ThermoFisher Scientific, Waltham, MA, USA). Protein bands were quantified by densitometry using ImageJ software (National Institute of Health). Levels of proteins of interest were normalized to GAPDH levels.

### Alizarin Red Staining and quantification

To assess the mineralization of cells after dexamethasone and calcium stimulation, an Alizarin Red staining was performed. Cells were fixed in 4% paraformaldehyde (PFA) solution diluted in phosphate buffered solution (PBS) for 15 min at RT and rinsed twice in PBS and once in distilled water. 1% Alizarin Red solution diluted in water and adjusted to pH 4.1-4.2 was added to cells and incubated for 30 min at RT under agitation. Cells were rinsed several times with distilled water and imaging was performed on an inverted Nikon Ti-S/L100 microscope, equipped with a Nikon DS-Ri2 camera using a CFI Plan Apochromat K 20×objective (NA: 0.75, WD: 1.0). Images were analyzed using NIS-Elements software (version 5.30.06, Nikon). Quantification of the staining was performed using 10% cetylpyridinium chloride (CPC) diluted in sodium phosphate buffer. CPC was incubated ON at RT under gentle agitation. CPC solution was transferred in a 96-well plate and absorbance was measured at 562 nm using a ClarioStar spectrophotometer (BMG Labtech, Champigny-sur-Marne, France). Concentration was determined using a standard curve of Alizarin Red diluted in CPC solution.

### Pull-down assay

HMSCs were transfected with a biotinylated mimic of miR-125b-5p or biotinylated negative control sequence. After 7 and 14 days cultured in basal medium, calcium-saturated medium or osteogenic medium, cells were harvested in lysis buffer (20 mM Tris-Base; pH 7.5, 100 mM KCl and 5 mM MgCl_2_) added with 0.05 % Tween 20, 50 U RNAse OUT (Invitrogen, Waltham, MA, USA) and proteinase inhibitor cocktail (PIC, Sigma, Saint-Louis, MO, USA). Previous to pull-down, 150 µl of Dynabeads M-270 streptavidin (Invitrogen, Waltham, MA, USA) were washed 3 times with lysis buffer. Beads were blocked during 2 h, under agitation at 4°C in 800 µl of lysis buffer added with 0.05% Tween 20, 50 U RNAse OUT, PIC, 200 yeast tRNA (1 mg/ml, Invitrogen, Waltham, MA, USA), and BSA 1%. After two washes, 150 µl of harvested cells were incubated with the beads for 4 h at 4°C under agitation. Beads were delicately washed 3 times in 500 µl of lysis buffer and resuspended in TRIzol LS (Invitrogen, Waltham, MA, USA) and extracted for RNAs following the manufacturer’s protocol.

### Statistics

Statistics were performed on Graph Pad Prism 10.0.2. All data are expressed as the mean ± standard deviation (SD). Experiments were performed at least 3 independent times (N=3). Normality was checked using the Shapiro-Wilk normality test. Significant differences (*=p < 0.05, **=p < 0.01, ***= p

< 0.001 and ****= p < 0.0001) between groups were assessed using a one-way ANOVA followed by a Tukey-Kramer post hoc ANOVA test. Nonparametric data were analyzed using a Mann-Whitney.

## Data availability statement

Upon publication, all data associated with this manuscript will be made publicly available or upon request from the corresponding author (V.Joris)

## Acknowledgments

This work is supported by the Province of Limburg, Limburg Invests in its Knowledge Economy (LINK). We would like to thank Maria Paula Marks, Maria Eischen-Loges and Steven Vermeulen from the MERLN institute for their technical help.

## Authors contribution

VJ: design of the work, data acquisition, data analysis, writing and review of manuscript ERB: contribution in design of the work, review of the manuscript.

MvG: conception and design of the project, data analysis, writing and review of the manuscript, fund raising

## Figure legends

**Figure S1:**
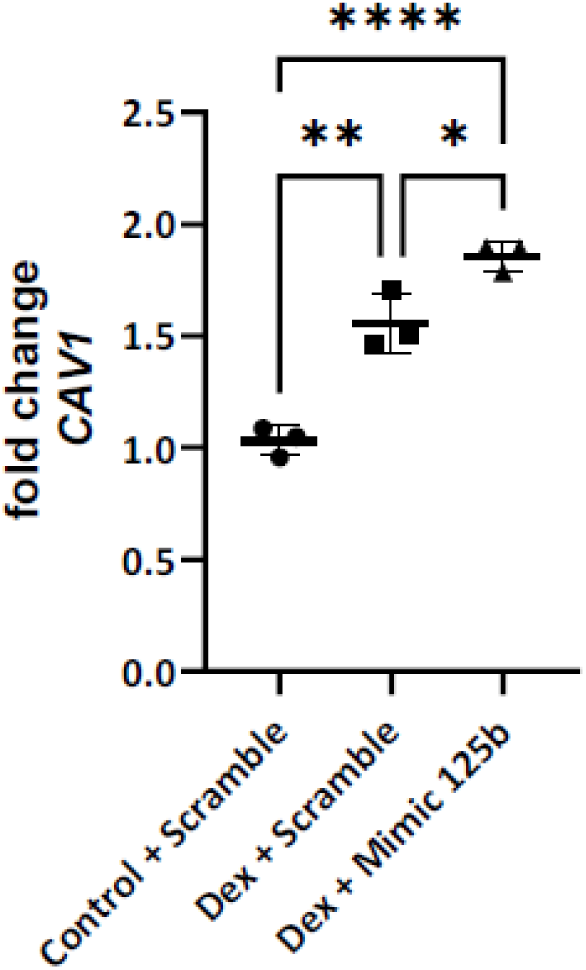
Dexamethasone and miR-125b mimic induced increase caveolin-1 (Cav1) expression. hMSCs were transfected with scramble or mimic and cultured In control dexamethasone medium. Cav1 expression was measured using qPCR. \**p* < 0.05; \*\**p* < 0.01; \*\*\*\**p* < 0.0001 based on fold change relative to control + scramble.

